# Modifications to the gut microbiome alter bone matrix proteomics and fracture toughness at the cellular scale

**DOI:** 10.1101/2025.11.21.689836

**Authors:** N.K. Hunt, C. Liu, J. Liu, Q. Tang, S.J. Stephen, B. Wang, C.A. Schurman, C.D. King, Q. Liu, D. Vashisth, B. Schilling, M. Hassani, C.J. Hernandez

**Author notes:** **Corresponding Author**: Christopher J. Hernandez. These authors contributed equally to this work (N.K. Hunt, C. Liu).

## Abstract

The gut microbiome can regulate the strength of bone matrix but the specific changes in matrix function and composition are not yet understood. Here we introduce micropillar splitting to determine the fracture toughness of matrix at the cellular scale in concert with proteomic analysis of bone matrix when the gut microbiome was altered by oral antibiotics (ampicillin+neomycin). Male mice were divided into four groups (n = 3-4/group): (1) Unaltered (no alteration), (2) Continuous (alteration from 4-24 weeks), (3) Delayed (alteration from 16-24 weeks) and (4) Reconstituted (alteration from 1-16 weeks following by reconstitution). Micropillars (5 µm diameter) were fabricated using focused ion beam milling on femur cross-sections in regions of matrix formed either before or after changes in the microbiome (16 weeks) (n = 4/group). Proteomics was used to identify differences in matrix protein composition. Micropillar fracture toughness differed by group (p < 0.001) and region (p < 0.001). Fracture toughness in the Unaltered group (1.34 ± 0.32 MPa√m, mean ± SD) was substantially greater than the Continuous group (0.95 ± 0.20) and the Delayed group (0.90 ± 0.21) but not different from the Reconstituted group (1.22 ± 0.25). Bone matrix formed from 16-24 weeks of age had lower fracture toughness than matrix formed before 16 weeks of age in all groups. Notably, micropillar splitting was substantially more precise than whole bone testing; whole bone notched 3-point bending tests did not detect differences in fracture toughness. Proteomics identified 46 extracellular matrix proteins that were differentially abundant between groups, including decreased abundance of Periostin (q < 0.001) and Emilin-1 (q < 0.001) in groups with impaired bone matrix. These findings demonstrate that modifications to the gut microbiome lead to changes in bone matrix throughout cortical bone volume and establish micropillar splitting as a high-precision approach for characterizing matrix material properties.

**Plain Language Summary:** This study investigated the effect of the gut microbiome on the brittleness of bone as a material. We used a new technique for measuring the brittleness of bone that involves making microscopic pillars on the bone surface and splitting them down the middle to measure a material property called fracture toughness. Mice with altered gut microbiomes had bone with reduced fracture toughness (by 35-40%). Restoration of an altered gut microbiome for two months reversed the effects. The effect of the microbiome on bone occurred throughout the whole bone and was not limited to regions of the bone formed after a change in the gut microbiome.

## 1.0 INTRODUCTION

The primary concern of bone disease is impairment of whole bone strength leading to fragility fracture. Therapies to address bone fragility, such as antiresorptive agents (e.g., bisphosphonates, denosumab), and anabolic agents (teriparatide, abaloparatide), work to maintain or even increase bone volume and bone mass by preventing bone resorption and/or increasing bone formation^1,2^. While successful, these agents do have limitations; after 3-5 years of treatment, gains in bone mineral density slow, and the risk of adverse side effects increases^3^. Alternative methods are needed to improve bone fragility and reduce fracture risk beyond what is currently possible. The strength of a whole bone is determined by bone mass and bone mineral density, whole bone morphology, and matrix composition. Bone matrix includes a mineral component, which primarily contributes to material strength and stiffness, as well as an organic component that primarily contributes to toughness ^4–10^.

Previous studies have shown that alterations to the composition of the gut microbiome can cause reductions in whole bone strength that are not explained by whole bone geometry, implicating changes in bone matrix^11^. Recently, we showed that modifications to the microbiome could be used to impair or improve the strength of bone matrix in a skeletally mature animal over time periods insufficient for matrix turnover by osteoclast/osteoblast activity. These findings suggest that alterations to the gut microbiome may influence all the bone matrix, not only the matrix formed after the microbiome is altered. However, assessment was limited to bending tests of whole bone, which provide only estimates of the material properties of the matrix ^12^. Additionally, differences in mineral composition at the microscale, measured using Raman spectroscopy (mineral:matrix ratio; crystallinity, etc.), could not explain the differences in mechanical performance. It is unclear what changes in matrix constituents may have caused the alterations in matrix performance.

The materials science community has developed several approaches to directly evaluate the mechanical properties of micro-scale samples that can be used to study bone from small animal models. Nanoindentation has been used to characterize bone material properties at the micrometer level for decades^13,14^. However, nanoindentation measurements are limited to hardness and stiffness, which are not failure properties and therefore do not describe how bone matrix breaks. More recently, a microscale testing method has been demonstrated that involves the creation of micropillars for materials testing using a focused ion beam (FIB). Zysset and colleagues pioneered the use of FIB to make micropillars to measure the compressive strength and Young’s modulus of bone matrix and demonstrated the approach in samples from large animals and humans^15^. While these studies have been insightful, compressive strength is only one influential failure property of bone matrix. Fracture toughness is another aspect of material failure that describes the brittleness of the material and is more sensitive to changes in collagens and other organic constituents of the matrix that contribute to fragility^16^. Fracture toughness has been assessed using cantilever samples prepared with FIB, although the approach is very time-intensive (seven hours of FIB preparation per sample)^17^. More recently, materials scientists have established ways of evaluating fracture toughness using micropillars, a technique called “micropillar splitting” ^18^.

There are several established methods for the characterization of the constituents of bone matrix. Spectroscopy (Raman, FTIR) has been used for decades to examine the chemical composition of bone matrix, providing measures including mineral to matrix ratio and mineral crystallinity^19–22^. Mineral content is a major determinant of matrix stiffness and strength (Young’s modulus and ultimate stress), but fracture toughness and brittleness are influenced more by the organic constituents of bone matrix. Although spectroscopy techniques can provide information about the composition or quality of the organic component of the matrix^23^, chemical characterization through mass spectrometry can provide more accurate assessments of the abundance of non-collagenous proteins and collagen cross-linking in matrix ^24^. Recently, proteomics approaches have been adapted to measuring bone matrix, making it possible to determine the abundance of thousands of different protein constituents in a bone sample along with post-translational modifications^5,24^. The specific changes in matrix protein composition associated with microbiome-induced changes in bone matrix are not known.

We asked the following research questions: (a) Do alterations to the gut microbiome affect matrix fracture toughness at the cellular scale?; (b) Does a change to the gut microbiome affect all matrix or only subsequently formed matrix?; and (c) what changes in bone matrix protein composition are induced by alterations to the gut microbiome? In this study, we developed and validated a micropillar splitting protocol for murine bone samples and applied this approach to determine changes in fracture toughness under conditions of an altered gut microbiota. We used micropillar splitting to evaluate microbiome-induced changes in bone fracture toughness and whole-bone proteomics to evaluate microbiome-induced changes to bone matrix protein composition. Additionally, we compared our results with whole-bone bending tests and Raman spectroscopy from the same cohort^11,25^. We hypothesize that gut microbiome-induced changes in bone matrix occur throughout the bone cross-section and are not limited to regions of bone formed after the change in the microbiome.

## 2.0 METHODS

### Animals

This study analyzes a subset of specimens from previously reported studies^11,25^. Here we briefly review the experimental plan. The Institutional Animal Care and Use Committee approved all experimental procedures. Male C57BL/6J mice were obtained from breeding pairs purchased from Jackson Laboratory (Bar Harbor, ME, USA). Only males were used because tissue strength was significantly different between treatment groups in males but not in females in the previously reported studies^11^. Upon weaning, offspring from breeding cages were distributed at random into cages, and treatment groups were randomly assigned to individual cages. Animals received housing in plastic cages containing ¼-inch corn cob bedding (The Andersons’ Lab Bedding, Maumee, OH, USA), standard laboratory chow (Teklad LM-485, Envigo Diets, Madison, WI, USA), water ad libitum, and cardboard refuge environmental enrichment huts (Ketchum Manufacturing, Brockville, Canada) under a 12:12 dark/light cycle. Until three weeks of age, mice remained housed with their dam in a specific pathogen-free facility. At three weeks of age, animals were randomly assigned to cages (4-5 animals/cage, 12-17 animals/group) separated by sex, and cages were subsequently transferred to a conventional housing facility. To minimize cage-to-cage variation in gut microbiota within experimental groups, soiled bedding was mixed weekly between cages of animals from the same experimental groups until 12 weeks of age.

### Study Design

This study examined samples from four groups with differences in bone matrix strength determined with three-point bending: Unaltered (normal bone matrix strength), Continuous (impaired bone matrix strength), Delayed (impaired bone matrix strength), and Reconstituted (normal bone matrix strength) (Figure 1A, B). The Unaltered group received standard drinking water throughout the experimental period. The dosed groups (Continuous, Reconstituted, and Delayed) received antibiotic treatment in drinking water consisting of 1 g/L ampicillin and 0.5 g/L neomycin. These antibiotics have low oral bioavailability and minimal uptake by the intestinal barrier, resulting in primary effects on gut microbiota composition with limited systemic distribution. Previous studies demonstrated that dosing with this antibiotic cocktail causes selective depletion of the gut microbiota (removes some but not all organisms) and leads to reduced bone matrix strength. The Continuous group received antibiotic dosing from weaning (4 weeks of age) until euthanasia (24 weeks of age). The Delayed group received standard drinking water until 16 weeks of age, followed by antibiotic-containing drinking water from 16-24 weeks of age. The Reconstituted group received antibiotics in drinking water from 4-16 weeks of age, followed by standard drinking water without antibiotics from 16-24 weeks to facilitate microbiota recovery. At 16 weeks, the Reconstituted group received fecal microbiota transplant (FMT) from sex- and age-matched Unaltered mice to repopulate the gut microbiome with Unaltered mouse gut microbiota. Fluorescent bone formation markers were administered at 12, 16, and 24 weeks of age to identify regions of bone matrix formed at different experimental timepoints. Modifications to the gut microbiota were confirmed with sequencing of fecal pellets (see^11^ for details). Animals underwent euthanasia at 24 weeks of age. At euthanasia, femurs were collected immediately, wrapped in phosphate-buffered saline (PBS)-soaked gauze and plastic wrap, and stored at −80°C. Micropillar and proteomics were performed on the distal portion of the femur collected after whole bone testing.

**Figure 1:**
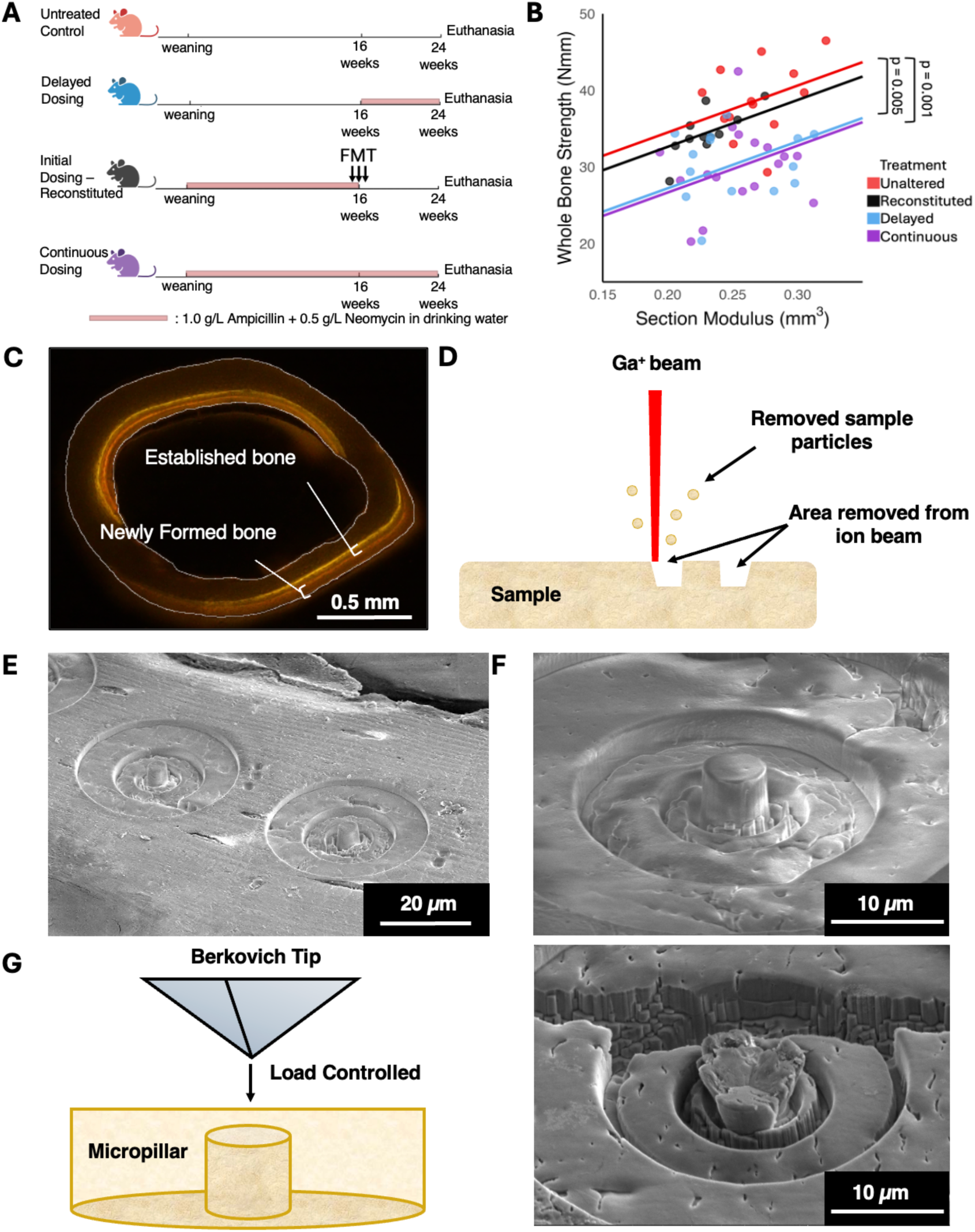
(A) Male C57Bl/6 mice were randomly divided into four experimental groups in which the gut microbiome was selectively depleted by antibiotics added to drinking water at different periods of life. (B) Graphical illustration of the ANCOVA analysis showing that the Delayed and Continuous groups have lower whole bone strength than expected from geometry (section modulus). Pairwise comparisons are indicated with p-values adjusted for multiple comparisons. Adapted/reprinted with permission from Liu, C., et al. 2024^11^. (C) The femur mid-diaphysis is shown with shaded regions indicating cortical bone formed during 4-16 weeks of age or 16-24 weeks of age, as indicated by bone formation labels. (D) Focused Ion Beam (FIB) lithography was used to ablate material on the surface of a cross-section of the bone, leaving a micropillar. (E, F) Scanning Electron Microscope (SEM) image of micropillars on the surface of bone samples before micropillar splitting was performed. (G) Micropillars were split using a Berkovich tip. (E) SEM image of a micropillar after micropillar splitting.

### Fecal Microbiota Transplant Preparation

Two fresh fecal pellets were collected from age- and sex-matched Unaltered mice (receiving only standard drinking water) and immediately transported in sterile 1.5 mL microfuge tubes to an anaerobic chamber. Fecal pellets were suspended in 1 mL of PBS containing 0.05% L-cysteine and allowed to soften for 15 minutes before undergoing vortexing for 3 minutes to achieve homogenization. To separate bacteria from fibrous material, samples were centrifuged for 5 minutes at 150 rpm. The supernatant was collected and pooled from multiple animals into a sterile bottle. The resulting fecal slurry was stored in an anaerobic jar and transported to the animal facility. Reconstituted group recipient mice underwent fasting for 3 hours prior to receiving 150 μL of donor fecal slurry via oral gavage. Mice were placed in fresh cages following gavage. Fresh fecal transplants were administered to each mouse daily for three consecutive days. Following FMT, fecal samples were collected every two weeks to verify gut microbiota recovery.

### Micropillar Sample Preparation

A subset of samples from each group was submitted to micropillar splitting (n = 4 samples from each group) to obtain fracture toughness. First, femora were dehydrated in 100% ethanol for 72 hours and embedded in poly methyl methacrylate (PMMA) using an accelerated PMMA embedding method to minimize PMMA infiltration into perilacunar spaces. PMMA-embedded femora were sectioned into approximately 1-mm-thick slices using a low-speed saw. Sections were mounted onto aluminum SEM stubs using Loctite 460 and dried overnight under a 500 g weight. Low-viscosity adhesive was essential to ensure rigid bonding between samples and SEM stubs. Samples underwent sequential polishing with diamond lapping films of 30, 9, 3, 1, and 0.1 microns. Polishing speeds were set at 60 rpm for 30- and 9-micron films, 45 rpm for 3-micron films, and 30 rpm for 1- and 0.1-micron films. Final polishing was performed using 0.05-micron water-free colloidal silica. Alcohol-based, water-free lubricants were applied throughout the process to prevent unwanted sample rehydration. Finally, samples were sputter-coated with a 3 nm thick gold/palladium layer to minimize charging during scanning electron microscopy imaging.

### Micropillar Manufacturing Using Focused Ion Beam

Micropillar positioning was selected based on fluorescent images of bone cross-sections. Bone formation markers applied to animals at 16 weeks of age were located on average 35 μm from the anterior-lateral periosteal surface. On each sample, two micropillars were fabricated within 20-30 μm of the periosteal edge to characterize bone matrix formed after 16 weeks of age (designated as “Newly Formed”), while an additional three micropillars were positioned within 40-50 μm from the periosteal surface to represent tissue formed before 16 weeks of age (designated as “Established”). Micropillars were fabricated on transverse femur sections using a ThermoFisher FEI Scios Dual-Beam Scanning electron microscope/Focused Ion Beam (SEM/FIB) (Thermo Fisher, Waltham, Massachusetts). After sample loading and vacuum achievement, scanning electron microscopy (SEM) at 5.0 kV and 1.6 nA was employed to identify regions of interest. Once regions were located, the focused ion beam was activated at a relatively low current (30 keV and 30.0 pA) to ensure proper SEM and FIB alignment and focusing on the desired sample locations. Validation studies using silica glass were prepared at Cornell University while hydrated v. dry bone samples and study specimens were prepared on the same model instrument at University of California, Davis.

Micropillars were fabricated through gradual material removal via milling of five concentric rings (Figure S1, Table S1). The first ring removal featured an outer diameter of 40 microns, an inner diameter of 20 microns, and a depth of 1 micron. This ring was milled with high current (7 nA) to rapidly remove bone matrix. The second ring, with an outer diameter of 22 microns and an inner diameter of 10 microns, had a depth of .51 microns and was milled using 5 nA current. The third ring, measuring 11 microns outer diameter and 6 microns inner diameter, had a depth of 1.5 microns using lower current of 3 nA. The fourth ring had an outer diameter of 7 microns and an inner diameter of 5.3 microns at a depth of 1.5 microns, using a current of 1 nA to minimize damage to the final micropillar. The final step involved micropillar polishing to a 5-micron diameter and 5-micron depth using a 0.5-nA current for smooth finishing. After manufacturing, micropillar diameter and height were determined by scanning electron microscopy to be 5.01 ± 0.12 μm (mean ± SD) and 5.33 ± 0.33 μm for an aspect ratio of 0.94 ± 0.06. Pillars were manufactured in two regions: Newly Formed bone matrix and Established bone matrix. The distance from the periosteal surface for the Newly Formed bone was 21.21 ± 4.04 μm, and for the Established bone was 43.78 ± 7.05 μm. Micropillar milling and characterization required 60 minutes per micropillar. A total of 53 micropillars were generated (3-4 micropillars/bone*16 bones).

### Micropillar Splitting

Micropillar splitting was performed using a nanoindenter (Hysitron Nanoindenters TI990, Bruker, CA) fitted with a Berkovich tip. To ensure hydration, samples were soaked in Hank’s Buffered Saline solution for 2 hours. The excess saline above the micropillar was gently removed with a Kimwipe to keep the sample surface accessible, leaving droplets within the well surrounding the micropillar. Micropillar locations were identified and pillar centers were located by scanning 15 × 15 mm² areas using the scanning probe microscopy (SPM) mode with the nanoindenter. Testing was performed under load-controlled conditions, maintaining a constant loading rate-to-load ratio of 0.05 s⁻¹. The Berkovich tip provides a sharp edge for initiating a sharp crack in the bone, rather than a machined notch, allowing for more consistent test conditions as it eliminates the need to align the machined notch with the indenter. Tests were manually terminated upon detecting sudden displacement shifts indicating mechanical failure, while maximum force was set at 10 mN. Time, indentation depth, and reaction force were recorded throughout experiments. The technique described herein was validated by testing of (1) silica glass and (2) bone in dry and hydrated bone conditions (see Supplementary methods for detail).

### Nanoindentation

Nanoindentation was performed on tissue adjacent to the micropillar using the same indenter. Six nanoindentations were performed in each region (Newly Formed v. Established) with a minimum spacing of 10 microns. A maximum load of 6 mN was applied at a constant loading rate of 0.355 mNs^-1^, followed by a holding time of 30 seconds. For validation studies evaluating the effects of hydration, dry samples were first tested with nanoindentation and micropillar splitting, then submerged in Hank’s Buffered Saline Solution, and subsequently tested again using nanoindentation and micropillar splitting on neighboring regions under hydrated conditions.

### Finite Element Modeling

Fracture toughness was calculated from experimental results using the approach established by Sebastiani et al.^18^. Fracture toughness is calculated as follows:

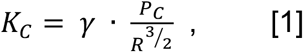

where P_C_ is instability load (mN), R is micropillar radius (µm) and γ is the micropillar splitting threshold, a constant determined from finite element models of the micropillar splitting experiment. A finite element model of the micropillar was created with material properties elastic modulus (E), Poisson’s ratio (ν), and cohesive zone fracture energy (G) (Figure S3, Table S3). The crack plane was simulated as a zero-thickness cohesive interface (constants in the simulation included traction-separation stiffness and maximum stress for damage initiation). The indentation of the Berkovich indenter was performed in a quasi-static, displacement-controlled manner. Gamma was determined from critical load (P_C_) at crack destabilization (Figure 2A). Six finite element models were performed to confirm that the value of gamma was not sensitive to micropillar aspect ratio (diameter to height) or the energy release rate in the cohesive elements^26^. We determined a pillar-splitting threshold value (γ) for the hydrated, Unaltered bone matrix of 0.94 (Table S3).

**Figure 2:**
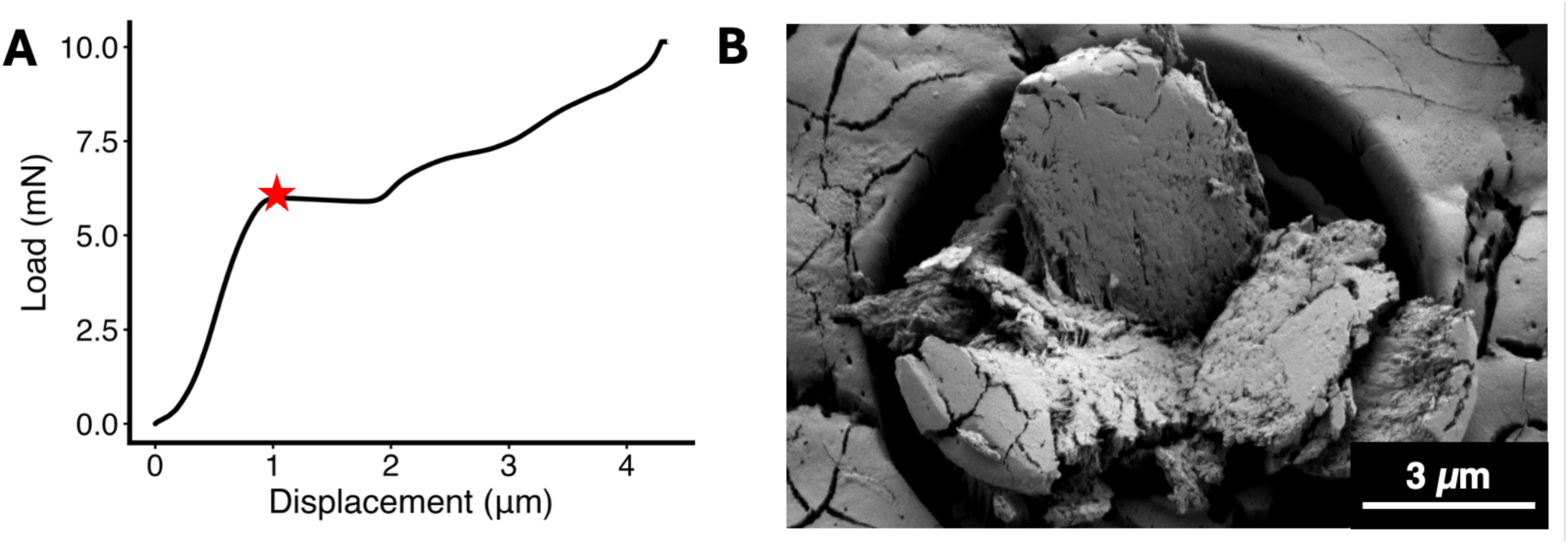
(A) Load versus displacement plot obtained from fully splitting a hydrated micropillar. The indenter was allowed to continue to displace through the full pillar, beyond the critical load point. The critical load of failure is marked with a red star. (B) An SEM image of a fully split micropillar.

### Micropillar Splitting Method Validation

For method validation, micropillars (n = 6) were fabricated on fused silica and tested using micropillar splitting with a nanoindenter within the SEM. Fused silica was selected because it is homogenous, previous studies had examined its behavior under micropillar splitting, and it was possible to compare results to fracture toughness in the literature. To evaluate the effects of hydration on micropillar splitting of bone, we compared the fracture toughness of micropillars from the same Unaltered samples in dry (n = 8) and hydrated (n = 5) conditions.

### Proteomics

The distal portion of the femur for all groups (n = 5/group) was collected after notched three-point bending and submitted to proteomic analysis by the Schilling Laboratory at the Buck Institute of Aging (Novato, CA) using previously established protocols^5,24^. Bone marrow was removed before samples were demineralized with 1 mL of 1.0M HCl overnight. Samples were frozen in liquid nitrogen and pulverized. Pulverized samples were transferred into 800 μL of extraction buffer in fresh Eppendorf tubes for 72 hours at 4°C. To separate bone matrix from supernatants containing soluble proteins, samples were centrifuged for 3 minutes at 15,000 ×g. The supernatant was filtered to remove guanidine hydrochloride by centrifuging through 3 kDa centrifugal filters at 12,000 ×g for 20 minutes and washed with 10 mM Tris-HCl. Isolated protein was resuspended in Tris-HCl and quantified using bicinchoninic acid assay. For protein digestion, the S-Trap mini spin column method was employed. Digested samples were resuspended in 0.2% formic acid, and indexed Retention Time Standards were added to samples per manufacturer’s instructions^27^. Mass spectrometric analysis was performed using reverse-phase HPLC-MS/MS in data-independent acquisition (DIA) mode^28–30^ using a Dionex UltiMate 3000 system and Orbitrap Exploris 480 mass spectrometer (ThermoFisher, San Jose, CA). DIA data was processed using Spectronaut using direct DIA for both protein and peptide levels. See supplementary mass spectrometry methods for a more detailed explanation of protein extraction, proteolytic digestion and desalting, mass spectrometry analysis, and data processing.

### Statistical Analysis

Differences among groups in whole bone fracture toughness and micropillar fracture toughness were evaluated using one-way analysis of variance (ANOVA). Pairwise comparisons were performed using the Tukey test to adjust for multiple comparisons. The effects of treatment and bone matrix region (Newly Formed or Established) on bone fracture toughness was determined using linear mixed-effects model using the lmer function from the lmerTest package in R (v3.1.3). The model included fixed effects for treatment (Unaltered, Delayed, Continuous, Reconstituted), region (Newly Formed or Established), and their interaction (treatment × region), with bone sample included as a random variable to adjust for multiple micropillars from each sample. Statistical significance was defined as p < 0.05. Analyses were conducted using R Version 2025.09.1+401.

Differential proteomics abundance analyses were performed pooling samples from groups with reduced bone matrix strength groups (Continuous and Delayed) and samples from groups with normal bone matrix strength groups (Unaltered and Reconstituted) using paired t-tests, and p values were corrected for multiple testing using the Storey method^31^. Proteins and post-translational modification sites were considered significantly altered if protein groups with at least 2 unique peptides had q values < 0.05 and absolute Log_2_(fold-change) > 0.58.

## 3.0 RESULTS

### Micropillar Testing and Nanoindentation

Fracture toughness was significantly different among treatment (p < 0.001) and region (p < 0.001, GLM) (Figure 3A). The fracture toughness obtained by hydrated micropillar splitting of the Unaltered group was 34.8% (pooled Newly Formed and Established regions: 1.34 ± 0.32 MPa√m) greater than the Continuous group (0.95 ± 0.20 MPa√m, p = 0.002) and 40.0% greater than the Delayed group (0.90 ± 0.21 MPa√m, p < 0.001) (Figure 3A). The Reconstituted group (1.22 ± 0.25 MPa√m) was not significantly different from the Unaltered group (p = 0.614) or Continuous group (p = 0.096) but was significantly greater than the Delayed group (p = 0.038).

**Figure 3:**
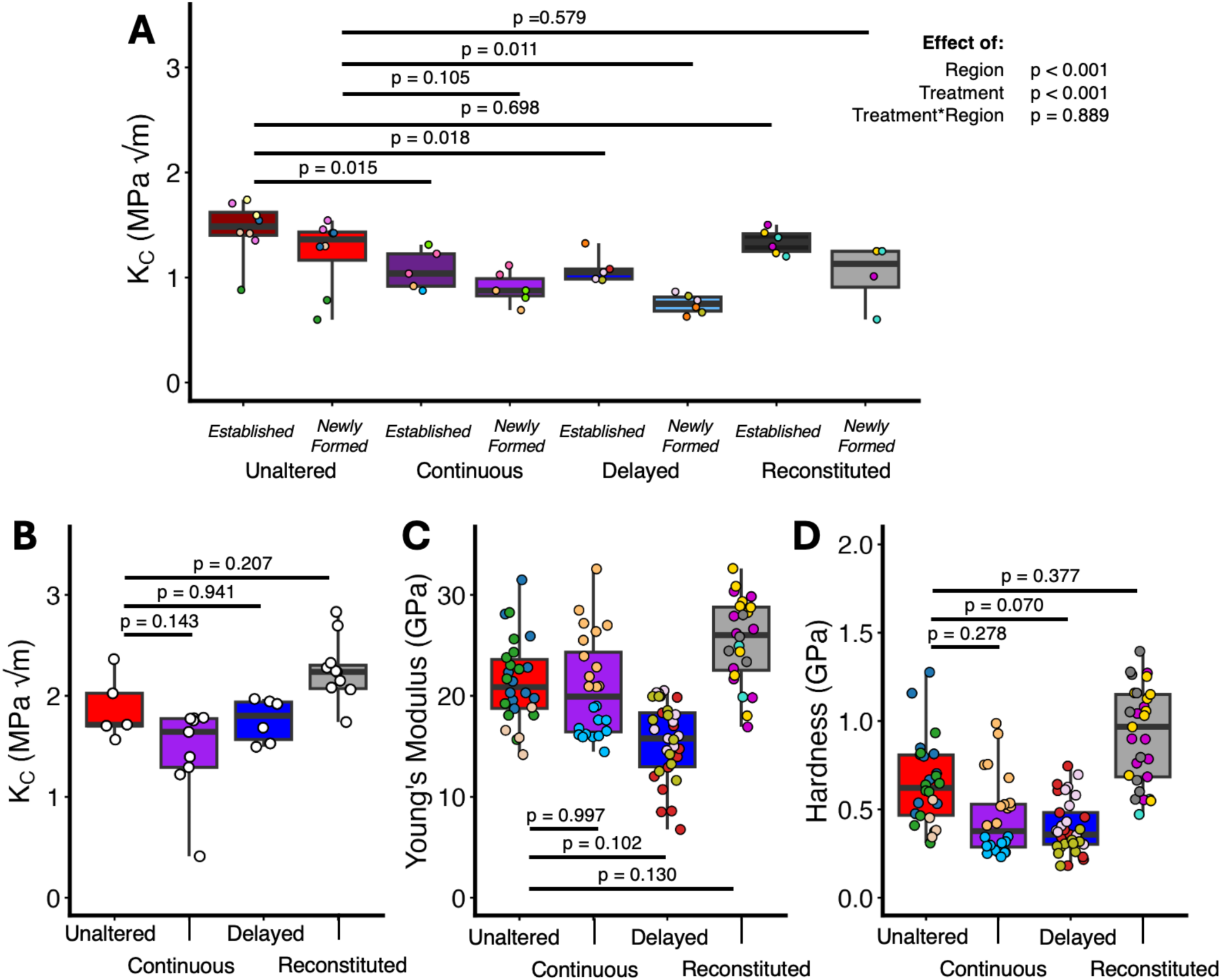
(A) Fracture toughness measured from hydrated micropillars in the Established (>40 µm from the periosteum) and Newly Formed (30-40 µm from the periosteum). Fracture toughness was significantly different among treatment and region. (B) Whole bone notched three-point bending results from a larger sample size of the same experimental cohort (as presented by Wang et al. 2025). No significant differences in whole bone fracture toughness were detected between groups. Nanoindentation measures of (C) Young’s modulus and (D) Hardness are shown. Young’s modulus and hardness are not significantly different among treatment groups. Data points within a group that are the same color indicate micropillars from the same bone cross-section

Fracture toughness in the matrix formed after 16 weeks of age (Newly Formed region) was lower than that in the matrix formed before 16 weeks of age (Established region) in each treatment group (p < 0.001). The interaction between treatment and region was not statistically significant (p = 0.889), indicating that the difference between Newly Formed and Established matrix regions did not differ across groups. Nanoindentation testing did not detect significant differences between Newly Formed and Established regions for Young’s modulus (p = 0.441) or hardness (p = 0.723) within any treatment groups. No significant differences in Young’s modulus were observed among treatment groups (Figure 3C). Similarly, no significant differences in hardness were detected among treatments (Figure 3D).

Hydration greatly influenced fracture toughness measures. The fracture toughness of dry micropillars was 51.4% lower than that of hydrated micropillars (Figure S2). Dry samples also had a significantly greater hardness (p = 0.006) and Young’s modulus (p = 0.008) as compared to the hydrated samples (Figure S2).

### Global Proteomic Analysis

Liquid chromatography-mass spectrometry proteomics analysis of the whole bone identified 1085 proteins with UniProKB-TrEMBL IDs, all with at least two unique peptides that were associated with the full protein. From the 1085 identified proteins, 46 were differentially abundant between animals with reduced matrix strength (Delayed and Continuous pooled) and those with normal bone strength (Unaltered and Reconstituted pooled). Of those, 37 proteins were differentially downregulated, and 9 proteins were differentially upregulated in the reduced bone matrix strength group versus the normal tissue strength group. Key core matrisome and matrisome-associated proteins, such as Elastin microfibril interface–located protein 1 (Emilin-1) and Periostin (Postn), were identified as significantly downregulated in the groups with reduced matrix strength compared to groups with normal matrix strength (q < 0.001 for both proteins) (Table S4). The core matrisome consists of the structural components of the extracellular matrix (ECM) including ECM collagens, glycoproteins, and proteoglycans^32^. The matrisome-associated proteins include ECM-affiliated proteins, ECM regulators, and secreted factors that bind to the core matrisome^32^. Pathway analysis demonstrated that glutamine synthesis was the most substantially downregulated pathway, along with other significantly affected metabolic pathways, including glycosaminoglycan (GAG) degradation and metabolism. Post-translational modification analysis detected 1028 hydroxyproline (HYP) sites on collagens, with 103 downregulated sites and 3 upregulated sites on the Continuous versus the Unaltered groups (Figure S4).

**Figure 4:**
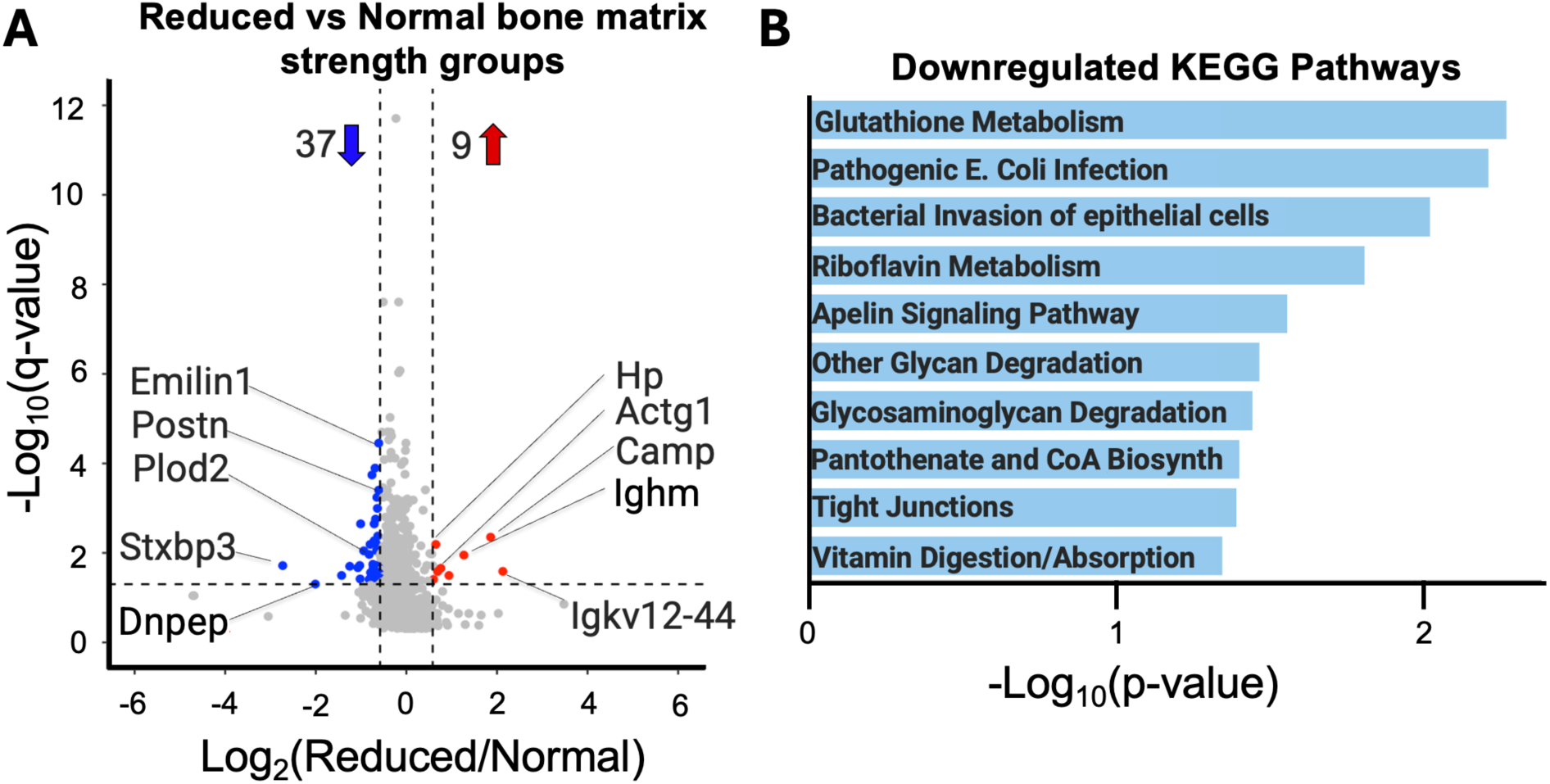
Proteomics of the whole bone provided a comparison of reduced bone matrix groups (Continuous and Delayed combined) to normal bone matrix groups (Unaltered and Reconstituted combined) (A) Volcano plot illustrating the 46 differentially abundant proteins of 1085 detected protein groups with 2+ unique peptides each. (B) Ingenuity Pathway Analysis showing the top ten downregulated KEGG pathways.

## 4.0 DISCUSSION

Alterations to the gut microbiome alter bone matrix fracture toughness in regions formed while the microbiome was altered as well as regions that were established before changes in the gut microbiome. These findings demonstrate that osteoclast/osteoblast mediated turnover of bone matrix is not necessary to enable microbiome-induced changes in bone matrix. The micropillar splitting technique shows high precision and sensitivity to differences between groups, detecting 34-40% reductions in fracture toughness among groups using only a small sample size (n = 4 animals/group). In contrast, traditional whole bone bending tests for fracture toughness in the same cohort did not detect differences between groups despite using a much larger sample size (n = 5-11/group). Proteomic analysis revealed downregulation of several matrix proteins in groups with reduced matrix strength, including Emilin-1 and Periostin, and also implicates alterations in metabolic pathways, including glutamine synthesis and glycosaminoglycan metabolism.

Interpreting our findings regarding differences between cellular scale measures of fracture toughness and whole bone bending measures of fracture toughness and strength requires a brief review of material properties. The strength of a structure (i.e. a whole bone) is influenced by material-level strength as well as material fracture toughness (see ^16^ for a conceptual review). For example, glass has high strength (hence it is used to make fiberglass) but low fracture toughness. A bone made entirely of glass would be prone to shattering due to low fracture toughness and would therefore display a low whole bone strength when subjected to bending, even though the material from which it is made has high strength. For this reason, the observed microbiome-induced reductions in fracture toughness at the cellular scale using micropillars are consistent with the reductions in tissue strength observed with whole bone bending (Figure 1B). Although micropillar splitting indicates differences in fracture toughness among groups, there were no significant differences in fracture toughness measured with notched three-point bending tests. We attribute this to the inherent limitations of notched bending test (described in detail in the original presentation of the method ^33^). Variability in fracture toughness measured by notched bending tests is caused by factors that are difficult to control: including variability in whole bone geometry along the length of the bone, the positioning and sharpening of the notch, and inherent heterogeneity of bone matrix, as results indicate, between the Newly Formed regions and Established regions. In contrast, micropillar splitting isolates material fracture toughness by testing standardized microscale specimens with negligible variance in geometry among micropillars, providing a more sensitive and direct assessment of matrix quality. The fracture toughness values obtained from micropillar splitting (0.90-1.34 MPa√m) are slightly lower than those typically reported from whole bone fracture toughness tests (human, longitudinal, 1.7 ± 0.3 MPa√m)^34^. We attribute the reduced fracture toughness to the fact that the micropillars are too small to include microstructures (cement lines, etc.) that provide extrinsic toughening mechanisms to resist crack growth, such as crack bridging and crack deflection^4^. We therefore speculate that the differences in fracture toughness among the mouse study groups in the current study were caused primarily by differences in intrinsic toughness mechanisms operating within the matrix.

Microbiome-induced changes in bone matrix quality compromise resistance to mechanical failure, independent of alterations in bone mass or architecture. The consistently lower fracture toughness in Newly Formed matrix compared to Established matrix suggests that mineralization and maturation state influence fracture resistance, consistent with previous reports that tissue mineral density and collagen cross-linking change with tissue age. Newly Formed bone matrix typically exhibits lower mineral content and reduced non-enzymatic collagen cross-linking compared to mature, Established bone. However, the lack of significant interaction between treatment and region indicate that alterations to the microbiome affects matrix already present in the skeleton and is not limited to matrix formed under conditions of a modified microbiome. Proteomic analysis revealed reduced abundance of extracellular matrisome-associated glycoprotein, Emilin-1, and matricellular protein, Periostin, in the groups with impaired matrix (Continuous and Delayed) compared to groups with normal matrix strength (Unaltered and Reconstituted). Emilin-1 is an extracellular matrix glycoprotein that regulates elastic fiber assembly and modulates mineralization in skeletal tissues. Periostin, a matricellular protein expressed by osteoblasts and osteocytes, plays crucial roles in collagen fibrillogenesis, matrix organization, and mechanotransduction. The downregulation of Emilin-1 and Periostin in groups with compromised matrix strength suggests disruption of matrix assembly and organization processes. Previous studies have demonstrated that Periostin-deficient mice exhibit reduced bone strength and altered collagen organization^35–37^. consistent with our observation that reduced Periostin correlates with diminished matrix material properties. In addition to the reduced abundance of extracellular matrix proteins, there was a notable reduction in hydroxyproline (HYP) modifications within collagen proteins. Of the 1028 HYP sites identified on collagens, 103 were downregulated in the impaired matrix strength group compared to the normal matrix group. HYP is a critical post-translational modification required for collagen triple helix stability and proper fibril formation^38,39^, and the downregulation of these sites may contribute to the observed matrix defects. While Periostin does not specifically bind to HYP-modified collagen sites, both are integral to collagen structure and stability, suggesting that their combined downregulation may exacerbate matrix disorganization. Furthermore, the reduction in several matrisome-associated proteins, including Annexin A7, a phospholipid-binding protein found in collagen-containing extracellular matrix and plasma membranes^40,41^, may further contribute to impaired collagen integrity. Annexin A7 is part of a family typically associated with binding with collagens and interacting with the membranes of matrix vesicles to determine vesicle shape and therefore influence hydroxyapatite crystal growth^42^. Pathway analysis of the differences in protein abundance associated with microbiome-induced impairment of bone strength includes the downregulation of several pathways. The most drastically downregulated pathway was glutamine synthesis, along with changes in glycosaminoglycan degradation and metabolism. Glutamine serves as a critical substrate for collagen synthesis and is essential for osteoblast and osteocyte function^43–48^.

Micropillar splitting offers several advantages over traditional whole bone mechanical testing for studies examining changes in matrix quality in mice. We observed a 34.8% difference between Unaltered and Continuous groups and a 40% difference between Unaltered and Delayed groups with micropillar splitting, while whole bone bending tests from the same cohort did not detect significant differences and observed a 22.8% difference between Unaltered and Continuous groups and only a 4.4% difference (not significantly different) between Unaltered and Delayed groups. Furthermore, the results of micropillar splitting tests show substantially less variance than whole bone bending tests. We attribute the greater precision of the micropillar testing to the fact that micropillars are not influenced of whole bone geometry, size, and structural organization. Reduced variability achieved through micropillar splitting enabled detection of significant differences with n = 4 animals per group in our study, compared to n = 12-15 typically required for whole bone mechanical testing in the literature.

The technical requirements for micropillar splitting are comparable to notched three-point bending in terms of time and labor. Furthermore, micropillars manufactured by FIB, have require less manual skill. In contrast, notched bending requires precise manual sharpening of the crack tip to ensure controlled crack propagation, a technique that introduces variability dependent on the operator. The ability to reduce sample size in studies using micropillars has the potential to reduce animal use and costs in animal husbandry, experimental treatments, and specimen preparation more than offsetting the costs and labor required for micropillar fabrication with FIB. Although micropillar fabrication requires dehydration and rehydration prior to testing, and whole bone testing does not, previous reports show that dehydration and rehydration have minimal impact on compact bone^49,50^.

A valuable feature of micropillar splitting is spatial resolution, which enables examination of specific matrix regions and cell-type-specific effects. In the present study, we distinguished between matrix formed before and after skeletal maturity to assess whether microbiome effects were limited to Newly Formed bone or also affect Established matrix. Spatial specificity cannot be achieved with whole bone mechanical testing, which averages properties across all tissue present. The ability to evaluate specific anatomical regions or tissue types makes micropillar splitting applicable to experimental questions currently addressed only through nanoindentation, including examination of periosteal versus endosteal bone, cortical versus trabecular tissue, and regions adjacent to metallic implants, regions within a fracture callus or focal pathologies where whole bone testing is not feasible.

The observation that alterations to the gut microbiome affect both Newly Formed and Established bone matrix has important mechanistic implications. That microbiome-induced changes in bone matrix were seen in Established bone suggests that the microbiome does not change bone through direct regulation of osteoblasts. We speculate that osteocyte-mediated perilacunar remodeling is a more likely mechanism. Perilacunar remodeling involves the modification of matrix composition, mineralization, and mechanical properties in the neighborhood of osteocytes and does not require osteoclast/osteoblast activity. Osteocytes respond to systemic metabolic signals, including changes in nutrient availability, hormone levels, and inflammatory mediators—all of which are influenced by the composition of the gut microbiome. While direct evidence of altered perilacunar remodeling in microbiome-altered mice awaits future investigation, the collective evidence points to osteocyte-mediated perilacunar remodeling as a plausible explanation for the observed widespread matrix changes.

In conclusion, the present study demonstrates that alterations to the gut microbiome reduce bone matrix fracture toughness throughout the entire bone matrix through mechanisms that extend beyond traditional bone modeling. Additionally, we demonstrate micropillar splitting as a means to provide a sensitive, precise, and spatially resolved method for assessing matrix quality with substantial advantages over conventional whole bone bending tests. The reversibility of matrix changes following microbiome reconstitution suggests that microbiome-targeted therapies could restore bone matrix quality even after prolonged disruption, hinting at the possibility of modulating bone matrix material properties in the adult skeleton.

## CONFLICT OF INTEREST STATEMENT

The authors declare no potential conflict of interest.

## ACKNOWLEDGEMENTS

Research reported in this article was supported by the US National Institutes of Health under award numbers R01AG067997, R21AR085253, R01DE019284 (CJH) and the US National Science Foundation (ENG-2125491, CJH). This work made use of the Cornell Center for Materials Research shared instrumentation facility and FIB acquisition supported by the NSF (DMR-2039380) as well as the UC Davis Center for Nano and Micro Manufacturing (CNM2), the UC Berkeley Materials Characterization Facility, and UC San Francisco Bioengineering & Biomaterials Correlative Imaging facility.

## DATA AVAILABILITY

The data underlying this article are available in the Dryad Digital Repository upon publication. Raw proteomics data and complete mass spectrometry datasets have been uploaded to the Mass Spectrometry Interactive Virtual Environment (MassIVE) repository, developed by the Center for Computational Mass Spectrometry at the University of California San Diego.

## Notes

### Competing Interest Statement

The authors have declared no competing interest.

