## Supplementary Materials for "Modifications to the gut microbiome alter bone matrix proteomics and fracture toughness at the cellular scale"

**Affiliations:**
[1]Department of Orthopaedic Surgery, University of California, San Francisco, CA, USA
[2]Department of Bioengineering and Therapeutic Sciences, University of California, San Francisco, CA
[3]Sibley School of Mechanical and Aerospace Engineering, Cornell University, Ithaca, NY
[4]Department of Biomedical Engineering, Center for Biotechnology and Interdisciplinary Studies, Rensselaer Polytechnic Institute, Troy, NY, USA.
[5]Shirley Ann Jackson Ph.D. Center for Biotechnology and Interdisciplinary Studies, Troy, NY, USA.
[6]Buck Institute for Research on Aging, Novato, CA, USA
[7]Rensselaer-Icahn School of Medicine at Mount Sinai Center for Engineering and Precision Medicine, Rensselaer Polytechnic Institute, New York, NY, USA.
[8]Materials Science and Engineering, Cornell University, Ithaca, NY
[9]Chan Zuckerberg Biohub San Francisco, CA, USA

*These authors contributed equally to this work

**Keywords**: bone biomechanics; fracture toughness; proteomics; microbiome; matrix

**Corresponding Author**: Christopher J. Hernandez

**Table of Contents:**

Supplementary Methods
Supplementary Results
Supplementary References

**Supplementary Methods**

*Micropillar Fabrication*

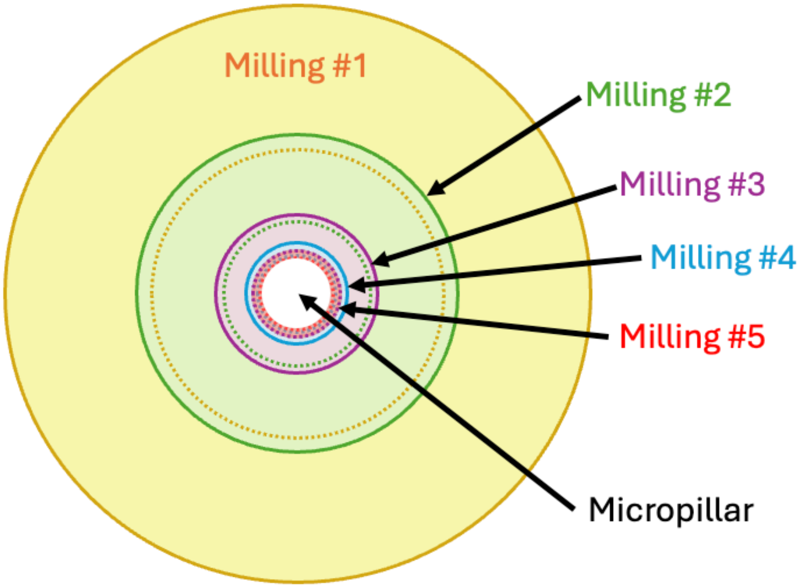


**Figure S1:** Micropillars were fabricated through gradual material removal via milling of five concentric rings that slightly overlap.

**Table S1**: The focused ion beam parameters for each milling round to manufacture the micropillars.

| Milling Round | Outer Diameter (μm) | Inner Diameter (μm) | Depth (μm) | Ion Beam current (nA) |
|---|---|---|---|---|
| 1 | 40 | 20 | 1 | 7.0 |
| 2 | 22 | 10 | 1.5 | 5.0 |
| 3 | 11 | 6 | 1.5 | 3.0 |
| 4 | 7 | 5.3 | 1.5 | 1.0 |
| 5 | 6 | 5 | 1.5 | 0.5 |

*Micropillar Splitting Method Validation*

For method validation, micropillars (n = 6) were fabricated on fused silica and tested using micropillar splitting with a nanoindenter within the SEM. Fused silica was selected because it is homogenous, previous studies had examined its behavior under micropillar splitting[1–3], and it was possible to compare results to fracture toughness in the literature. To evaluate the effects of hydration on micropillar splitting of bone, we compared the fracture toughness of micropillars from the same Unaltered samples in dry (n = 8) and hydrated (n = 5) conditions.

*Protein Extraction*

Femurs (n=5 per group) were cleaned of soft tissue and the periosteum before the epiphyses were removed at the growth plate and bone marrow removed by centrifugation to isolate the cortical diaphysis. Bones were wrapped in gauze moistened with Hank's Balanced Salt Solution (Corning #MT21022CV) supplemented with Complete Mini EDTA-free Protease Inhibitor (Roche #1183617000) and stored at −80°C until use for Data-Independent Acquisition (DIA) Mass Spectrometry (MS) analysis as previously described[4]. To remove residual soft tissue, thawed bones were incubated in a 1:1 mixture of Collagenase I (Millipore #SRC103) and Collagenase II (Worthington #LS004177) at 1 µg/µL for 5 minutes at 37°C. Samples then underwent three 10-minute rounds of sonication in fresh water. Bones were subsequently demineralized overnight in 1 mL of 1.2 M HCl (Fisher #A144) at 4°C[4,5]. The demineralized bone was flash-frozen in liquid nitrogen and pulverized using a Covaris CP02 cryoPREP automated dry pulverizer. The resulting powder was extracted in 800 µL of buffer containing 6 M guanidine hydrochloride (Sigma #G4505), 10 mM Tris-HCl (Sigma-Aldrich #252859), and 50 mM EDTA (Sigma-Aldrich #E4884) under rotation at 4°C for 72 hours[6]. After centrifugation (15,000 × g, 3 min), supernatants were collected and buffer-exchanged three times with 500 µL of 10 mM Tris-HCl (pH 7.0) using Amicon 3 kDa centrifugal filters (MilliporeSigma #UFC900308) at 12,000 × g for 20 minutes each. Samples were concentrated to 20 µL for downstream processing.

*Proteolytic Digestion and Desalting*

Protein concentrations were quantified using a BCA assay (Thermo Fisher #23227). 50 µg of soluble bone protein was diluted in 4% SDS and 50 mM TEAB. Reduction was performed with 20 mM dithiothreitol (DTT, Sigma-Aldrich #D9779) for 10 minutes at 50°C followed by 10 minutes at room temperature. Alkylation was then carried out with 40 mM iodoacetamide (IAA, Sigma-Aldrich #I1149) for 30 minutes in the dark at room temperature. Samples were acidified to 1.2% phosphoric acid (Sigma-Aldrich #79622) and diluted sevenfold in S-Trap binding buffer (90% methanol in 100 mM TEAB, pH ~7; Fisher #A452-1). Proteins were loaded onto S-Trap micro spin columns (Protifi #C02-micro-80) and centrifuged at 4,000 × g for 10 seconds. Columns were washed twice with binding buffer, followed by digestion with Trypsin (Promega #V5111) a 1:25 ratio, incubated for 1 hour at 47°C, and again overnight at 37°C. Peptides were eluted sequentially with 50 mM TEAB, 0.5% formic acid (FA; Honeywell #F0507) in water, and 50% acetonitrile (ACN; Honeywell #34851) in 0.5% FA, centrifuging at 1,000–4,000 × g as appropriate. After vacuum drying, samples were reconstituted in 0.2% FA, desalted using Oasis 30-mg cartridges (Waters #WAT094225), dried again, and resuspended in 0.2% FA at 1 µg/µL. Indexed retention time (iRT) standards (Biognosys #1900615) were added per the manufacturer's instructions[7].

*Mass Spectrometry Analysis*

LC–MS/MS of digested bone lysates was conducted using a Dionex UltiMate 3000 system coupled to an Orbitrap Exploris 480 (Thermo Fisher Scientific). Solvent A was 2% ACN/0.1% FA in water; solvent B was 80% ACN/0.1% FA. Peptides (400 ng) were first loaded on an Acclaim PepMap 100 C18 trap column (0.1 × 20 mm, 5 µm; Thermo #164535) at 5 µL/min for 5 minutes and then separated on an Acclaim PepMap 100 C18 analytical column (75 µm × 50 cm, 3 µm; Thermo #164570) at 300 nL/min. The elution gradient ranged from 2.5% to 39.2% solvent B over 165 minutes, ramping to 98% for 1 minute before re-equilibration. All samples were acquired in data-independent acquisition (DIA) mode[8–10]. Full MS spectra were collected at 120,000 resolution (AGC target: 3e6 ions, maximum injection time: 60 ms, 350-1,650 m/z), and MS2 spectra at 30,000 resolution (AGC target: 3e6 ions, maximum injection time: Auto, NCE: 30, fixed first mass 200 m/z). The DIA precursor ion isolation scheme consisted of 26 variable windows covering the 350-1,650 m/z mass range with an overlap of 1 m/z (Table S2) [10].

*Data Processing and Statistical Analysis*

Raw DIA data were processed using Spectronaut (v17.6.230428.55965). Cortical bone data were analyzed using a custom spectral library built from >150 independent mouse bone runs (DIA and DDA), comprising 71,206 peptides. Search parameters included trypsin/P digestion (allowing two missed cleavages), fixed carbamidomethylation on cysteines, and variable methionine oxidation and N-terminal acetylation. Dynamic, non-linear iRT calibration with precision profiling was applied. Protein identification was filtered at 1% FDR for precursor and protein levels. Quantification was based on the extracted ion chromatogram (XIC) areas of 3–6 MS2 fragments, with local normalization and sparse data filtering. For differential protein expression, unpaired t-tests were performed, and p-values adjusted for multiple testing using the Storey method[11–13]. Proteins were considered significantly changed if they met the following criteria: ≥2 unique peptides, q-value <0.05, and $|\log_2(\text{fold-change})| > 0.58$.

**Table S2**: Isolation scheme of the DIA acquisition method on the Orbitrap Exploris 480 mass spectrometer. The DIA precursor ion isolation scheme consisted of 26 variable windows covering the 350-1,650 m/z mass range with an overlap of 1 m/z.

| Window | Start m/z | Stop m/z | Center m/z | z | Isolation Width |
|---|---|---|---|---|---|
| 1 | 350 | 383 | 366.5 | 3 | 33 |
| 2 | 382 | 408 | 395 | 3 | 26 |
| 3 | 407 | 429 | 418 | 3 | 22 |
| 4 | 428 | 448 | 438 | 3 | 20 |
| 5 | 447 | 467 | 457 | 3 | 20 |
| 6 | 466 | 484 | 475 | 3 | 18 |

| | | | | | |
|---|---|---|---|---|---|
| 7 | 483 | 503 | 493 | 3 | 20 |
| 8 | 502 | 521 | 511.5 | 3 | 19 |
| 9 | 520 | 539 | 529.5 | 3 | 19 |
| 10 | 538 | 557 | 547.5 | 3 | 19 |
| 11 | 556 | 575 | 565.5 | 3 | 19 |
| 12 | 574 | 594 | 584 | 3 | 20 |
| 13 | 593 | 614 | 603.5 | 3 | 21 |
| 14 | 613 | 634 | 623.5 | 3 | 21 |
| 15 | 633 | 656 | 644.5 | 3 | 23 |
| 16 | 655 | 678 | 666.5 | 3 | 23 |
| 17 | 677 | 701 | 689 | 3 | 24 |
| 18 | 700 | 726 | 713 | 3 | 26 |
| 19 | 725 | 756 | 740.5 | 3 | 31 |
| 20 | 755 | 787 | 771 | 3 | 32 |
| 21 | 786 | 823 | 804.5 | 3 | 37 |
| 22 | 822 | 862 | 842 | 3 | 40 |
| 23 | 861 | 914 | 887.5 | 3 | 53 |
| 24 | 913 | 979 | 946 | 3 | 66 |
| 25 | 978 | 1077 | 1027.5 | 3 | 99 |
| 26 | 1076 | 1650 | 1363 | 3 | 574 |

### Supplementary Results

*Micropillar Splitting Validation*

Fused silica was chosen as a validation of micropillar splitting testing because it is a homogeneous material and has been previously characterized with micropillar splitting. We found our fracture toughness of fused silica tested within an SEM with active imaging (1.03 ± 0.21 MPa√m) to be slightly greater than the reported range of 0.58-0.78 MPa√m observed with macroscale testing (Figure S2D)[1–3,14]. The disagreement between micropillar splitting and the theoretical range could be due to the micropillar splitting experiments being conducted inside the SEM. Previous studies found micropillar splitting of fused silica in an SEM environment resulted in instability loads significantly larger than those observed with the conventional nanoindenter under ambient conditions[1].

Hydration of the micropillars greatly influenced Young’s modulus, hardness, and fracture toughness measures. Dry samples had a significantly greater hardness ($p = 0.006$) and Young’s modulus ($p = 0.008$) as compared to the hydrated samples (Figure S2A, B) but the fracture toughness was 51.4% lower than that of hydrated micropillars (Figure S2C).

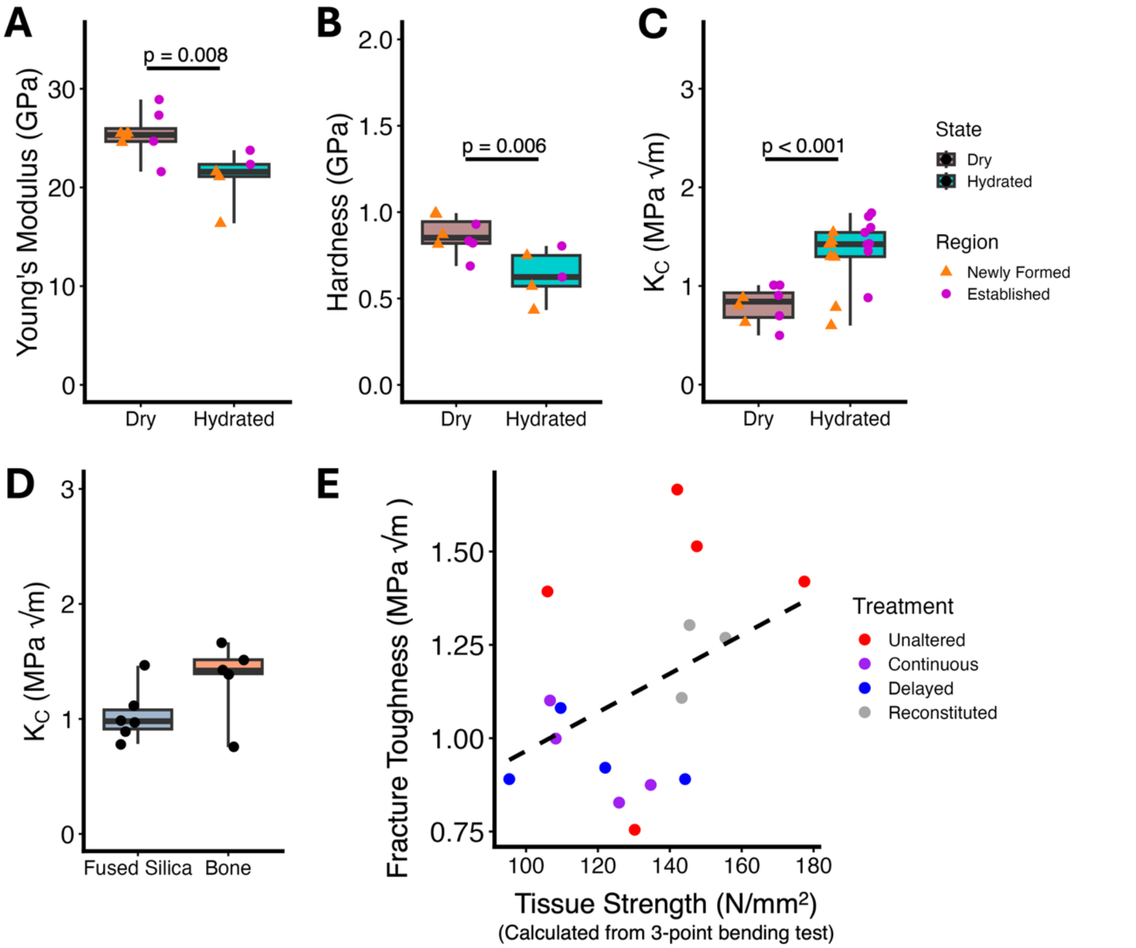


**Figure S2:** Nanoindentation of dry versus hydrated bone from the Unaltered treatment group provided (A) Young's modulus, (B) Hardness and (C) fracture toughness. (D) Micropillar splitting applied to fused silica provided a fracture toughness (1.03 ± 0.21 MPa√m). (E) Tissue strength calculated from 3-point bending versus fracture toughness calculated from micropillar splitting is shown ($R^2$ = 0.17).

*Finite Element Modeling*

We determined the micropillar splitting threshold value (γ) using finite element modeling, as described by Sebastiani et al. Simulations were performed with ABAQUS/CAE 2019 (Simulia, Providence, RI, USA). The three-dimensional model simulated contact between a rigid Berkovich pyramidal indenter (centerline-to-face angle of 65.3°) and a cylindrical micropillar of mouse bone with a 1:1 aspect ratio (height to diameter). The micropillar was modeled as an isotropic, elastic-perfectly plastic, and brittle material. To reduce computational cost, we employed a symmetry model that included one-third of both the indenter and pillar geometry (Figure S3A).

Symmetric boundary conditions were applied at the pillar side walls, while the base of the pillar was held fixed.

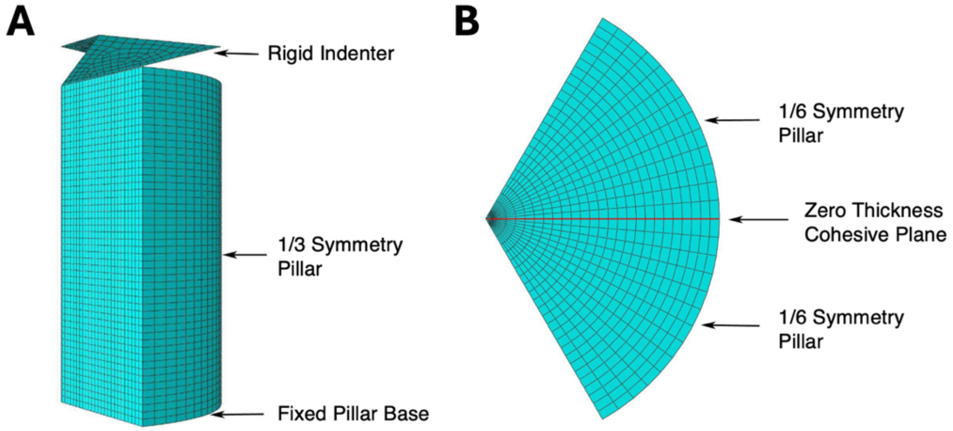


**Figure S3:** The finite element model is shown. The model uses ⅓ symmetry pillar splitting finite element model with a fixed pillar base, a cohesive plane in between, and a rigid indenter at (A) side view (B) top view.

**Table S3**: The modeling parameters and results for finite element analysis are shown for each treatment group. Critical load ($P_C$) was obtained from micropillar splitting.

| | **Unaltered** | **Continuous** | **Delayed** | **Reconstituted** | **Dry** |
|---|---|---|---|---|---|
| Young's Modulus (E) | 21 | 21 | 16 | 22 | 28 |
| E/Y | 52.5 | 52.5 | 40 | 55 | 70 |
| Poisson's Ratio | 0.35 | 0.35 | 0.35 | 0.35 | 0.3 |
| $P_C$ (mN) | 5.65 | 4.00 | 3.39 | 5.06 | 4.48 |
| γ | 0.94 | 0.94 | 1.05 | 0.95 | 0.71 |

*Proteomics*

**Table S4**: Proteomics identified 1060 proteins total with 2+ unique peptides. The table lists the 46 differentially abundant proteins between animals with reduced matrix strength (Delayed and Continuous) and those with normal bone strength (Unaltered and Reconstituted). Of those, 37 proteins were differentially downregulated (negative log fold change), and 9 proteins were differentially upregulated (positive log fold change) in the reduced bone matrix strength group versus the normal tissue strength group. Proteins are classified as core matrisome (ECM collagens, glycoprotein, proteoglycans), matrisome-associated (ECM-affiliated protein, ECM regulator, secreted factor), or non-matrisome.

| Gene | Detailed Name | Matrisome Classification | Log Fold Change |
|---|---|---|---|
| Stxbp3 | Syntaxin-binding protein | Non-matrisome | -2.73 |
| Dnpep | Dipeptidase | Non-matrisome | -2.01 |
| Mars | Methionyl-tRNA synthetase | Non-matrisome | -1.43 |
| Slc39a10 | Zinc transporter | Non-matrisome | -1.25 |
| Slc9a3r1 | Sodium/hydrogen exchanger | Non-matrisome | -1.07 |
| Pafah1b2 | Platelet-activating factor acetylhydrolase | Non-matrisome | -1.03 |
| Snx6 | Sorting nexin | Non-matrisome | -1.02 |
| Enpp1 | Ectonucleotide pyrophosphatase | Non-matrisome | -1.01 |
| Cope | Coatomer protein epsilon | Non-matrisome | -0.94 |
| **Plod2** | **Procollagen-lysine dioxygenase** | **ECM-regulator** | **-0.82** |
| Bzw1 | Basic leucine zipper transcription factor | Non-matrisome | -0.82 |
| Tcn2 | Transcobalamin II | Non-matrisome | -0.8 |
| **C1qtnf3** | **Complement C1q TNF-related protein** | **ECM-affiliated** | **-0.79** |
| Esd | Esterase D | Non-matrisome | -0.77 |
| **Emilin1** | **Elastin microfibril interface protein** | **ECM glycoprotein** | **-0.76** |
| Myo1d | Myosin ID | Non-matrisome | -0.76 |
| Hexb | Hexosaminidase B | Non-matrisome | -0.74 |
| **Ptn** | **Pleiotrophin** | **ECM glycoprotein/secreted factor** | **-0.73** |
| Lipc | Hepatic lipase | Non-matrisome | -0.71 |
| Arpc1b | Actin-related protein complex | Non-matrisome | -0.71 |
| Gdi1 | GDP dissociation inhibitor | Non-matrisome | -0.69 |
| Cnpy4 | Canopy homolog 4 | Non-matrisome | -0.69 |
| Idh2 | Isocitrate dehydrogenase 2 | Non-matrisome | -0.68 |
| Cisd1 | CDGSH iron sulfur domain protein | Non-matrisome | -0.68 |
| Fhl1 | Four and a half LIM domains 1 | Non-matrisome | -0.67 |
| Ckap4 | Cytoskeleton-associated protein 4 | Non-matrisome | -0.66 |
| Srsf5 | Serine/arginine-rich splicing factor 5 | Non-matrisome | -0.66 |
| Cdh1 | E-cadherin | Non-matrisome | -0.65 |
| **Postn** | **Periostin** | **ECM glycoprotein** | **-0.64** |
| Pls3 | Plastin 3 | Non-matrisome | -0.63 |
| **Anxa7** | **Annexin A7** | **ECM-affiliated protein** | **-0.61** |
| Gstm1 | Glutathione S-transferase M1 | Non-matrisome | -0.61 |
| Ranbp3 | Ran-binding protein 3 | Non-matrisome | -0.6 |
| Ube2h | Ubiquitin-conjugating enzyme | Non-matrisome | -0.6 |
| Gnb1 | G-protein beta 1 | Non-matrisome | -0.58 |
| Krt13 | Keratin 13 | Non-matrisome | -0.58 |
| Rab14 | Ras-related protein | Non-matrisome | -0.58 |
| Rps28 | Ribosomal protein S28 | Non-matrisome | 0.58 |
| Tf | Transferrin | Non-matrisome | 0.6 |
| Hp | Haptoglobin | Non-matrisome | 0.65 |

| | | | |
|---|---|---|---|
| Pzp | Pregnancy zone protein | Non-matrisome | 0.69 |
| Actg1 | Gamma actin | Non-matrisome | 0.76 |
| Mug1 | Murinoglobulin 1 | Non-matrisome | 0.94 |
| Ighm | Immunoglobulin heavy chain mu | Non-matrisome | 1.27 |
| Camp | Cathelicidin antimicrobial peptide | Non-matrisome | 1.86 |
| Igkv12-44 | Immunoglobulin kappa variable chain | Non-matrisome | 2.13 |

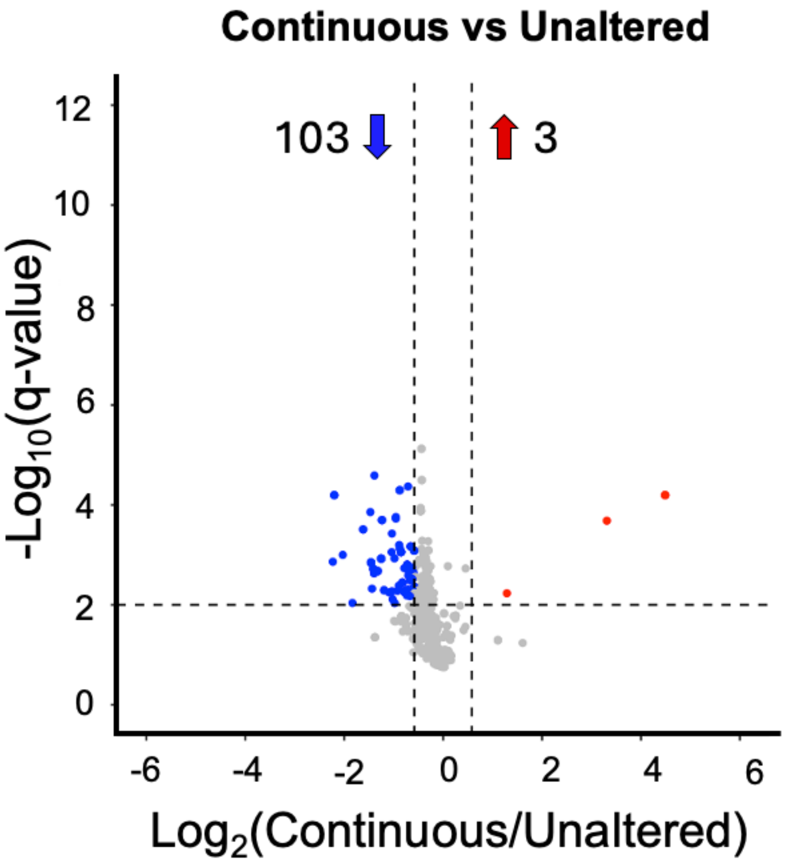


**Figure S4:** The Continuous group versus the Unaltered group hydroxyproline (HYP) site regulation. HYP site regulation: Q < 0.01 and Log2FC ≥ 0.58 (amount normalized to protein content). Of the 1028 total HYP sites detected, 103 were downregulated and 3 upregulated in the Continuous group versus the Unaltered group.
